# Differential requirement for IRGM proteins during tuberculosis infection in mice

**DOI:** 10.1101/2022.11.09.515519

**Authors:** Kaley M. Wilburn, Rachel K. Meade, Emma M. Heckenberg, Jacob Dockterman, Jörn Coers, Christopher M. Sassetti, Andrew J. Olive, Clare M. Smith

**Affiliations:** Department of Molecular Genetics and Microbiology, Duke University, Durham, United States; University Program in Genetics and Genomics, Duke University, Durham, United States; Department of Immunology, Duke University Medical Center, Durham, North Carolina, USA; Department of Microbiology and Physiological Systems, University of Massachusetts Medical School, Worcester, United States; Department of Microbiology and Molecular Genetics, College of Osteopathic Medicine, Michigan State University, East Lansing MI

## Abstract

*Mycobacterium tuberculosis* (*Mtb*) is a bacterium that exclusively resides in human hosts and remains a dominant cause of morbidity and mortality among infectious diseases worldwide. Host protection against *Mtb* infection is dependent on the function of immunity-related GTPase clade M (IRGM) proteins. Polymorphisms in human *IRGM* associate with altered susceptibility to mycobacterial disease, and human IRGM promotes the delivery of *Mtb* into degradative autolysosomes. Among the three murine IRGM orthologs, *Irgm1* has been singled out as essential for host protection during *Mtb* infections in cultured macrophages and *in vivo*. However, whether the paralogous murine *Irgm* genes, *Irgm2* and *Irgm3*, play roles in host defense against *Mtb* or exhibit functional relationships with *Irgm1* during *Mtb* infection remains undetermined. Here, we report that *Irgm1*^-/-^ mice are indeed acutely susceptible to aerosol infection with *Mtb*, yet the additional deletion of the paralogous *Irgm3* gene restores protective immunity to *Mtb* infections in *Irgm1*-deficient animals. Mice lacking all three *Irgm* genes (pan*Irgm*^-/-^) are characterized by shifted lung cytokine profiles at 4 and 24 weeks post infection, but control disease until the very late stages of the infection, when pan*Irgm*^-/-^ mice display increased mortality compared to wild type mice. Collectively, our data demonstrate that disruptions in the balance between *Irgm* isoforms is more detrimental to the *Mtb*-infected host than total loss of *Irgm*-mediated host defense, a concept that also needs to be considered in the context of human *Mtb* susceptibility linked to *IRGM* polymorphisms.

## INTRODUCTION

Interest in cell-autonomous immune mechanisms that act downstream of interferon (IFN) signaling led to the discovery of four IFN responsive families of dynamin-like GTPase proteins (1). Among these, the immunity-related GTPases (IRGs) have been implicated in host resistance to many intracellular pathogens, including *Toxoplasma gondii, Listeria monocytogenes, Mycobacterium tuberculosis (Mtb), Salmonella typhimurium*, and *Chlamydia trachomatis* in mouse models of infection (2). C57BL/6 mice possess 21 *IRG* genes, which are divided into two sub-classes (“GMS” or “GKS”) based on the amino acid sequence encoded in the G1 motif of their N-terminal GTP-binding domains (3). The genome of C57BL/6 mice contains three GMS genes (*Irgm1, Irgm2*, and *Irgm3*). By contrast, through a series of fascinating evolutionary events, *IRG* genes have mostly been lost from the human genome, leaving *IRGM* as the only known homologue of the murine GMS genes. It is expressed as five different splice variants (*IRGM*a-e) (3). Despite its high degree of similarity, *IRGM* was initially presumed to be a pseudogene due to its truncated GTP-binding domain and lack of IFN-dependent expression (4). However, subsequent studies have identified protective functions for *IRGM* in autoimmunity or immune responses to infection via its intersection with the autophagy pathway (5–9). What factors have driven the differential expansion and deletion of *IRG* genes between mice and humans, and what the relative fitness costs or benefits of retaining or losing the IRG system remain intriguing questions (10). Studies that expand our understanding of how the IRG system functions in mice during infection with diverse pathogens simultaneously offer useful points of comparison for examining the function of IRGM in humans.

*Mtb* is a facultative intracellular bacterium that causes the death of ~1.4 million people worldwide annually (11). *Mtb* has co-evolved with humans for thousands of years and is adept at manipulating the immune responses of macrophages, its primary host cell niche (12, 13). Multiple studies have proposed that *Irgm1* is important for host protection during mycobacterial infection. It was previously shown that mice lacking *Irgm1* exhibit extensive lung damage associated with large lesions, are unable to control *Mtb* burden, and rapidly succumb to aerosol infection (14). Similarly, *Irgm1*^-/-^ mice intravenously infected with either *Mtb* or *Mycobacterium avium* survive through the acute stage of infection but cannot successfully control bacterial growth and die by ~8-16 weeks post-infection (14, 15). The human ortholog, *IRGM*, has also been linked to control of mycobacterial infection. Various polymorphisms in *IRGM* are associated with increased or decreased risk of active pulmonary TB; however, these associations may be host population- and bacterial strain-dependent (16–21). Interestingly, in a cohort of the Han population of Hubei Province, China, there was a direct relationship between a variant haplotype (−1208A/-1161C/-947T) that decreased transcriptional activity of the IRGM promoter, reduced *IRGM* expression in patient PMBCs, and increased risk of pulmonary TB disease (19). Conversely, a haplotype (−1208A/-1161C/-947C) that increases *IRGM* transcription was associated with reduced TB disease risk in two Chinese cohorts (17, 19). Proposed explanations for the requirement of *Irgm1/IRGM* during mycobacterial infection include promotion of optimal macrophage phagolysosome function via autophagy, or prevention of IFNγ-dependent death of T cells that results in severe lymphopenia (5, 14, 15, 22).

Observations across studies that examined mice with additional *Irgm* deficiencies indicate that the three genes (*Irgm1, Irgm2*, and *Irgm3*) have non-redundant functions and complex inter-regulatory relationships, as evidenced by mice displaying differential susceptibilities to infection with various intracellular pathogens depending on which *Irgm* genes are inactivated (1). For example, *Irgm1*^-/-^ mice exhibit dysregulated host protection during infection with *S. typhimurium*, but this phenotype can be partially or entirely countered when mice are deficient in both *Irgm1* and *Irgm3* (*Irgm1/3*^-/-^) (23). Mice infected with *T. gondii* or *C. trachomatis* on the other hand require *Irgm3* expression for host protection (24). Although *Irgm*-deficient mice are universally susceptible to *T. gondii, Irgm2* and *Irgm1/3* have differential roles in the cell-autonomous response to infection, regulating the recruitment of distinct effectors to the parasitophorous vacuole (2, 25). However, whether *Irgm2* and *Irgm3* exhibit functional interactions with *Irgm1* that significantly influence the host response during mycobacterial infection is not yet established. In this work, we investigated if mice deficient in both *Irgm1* and *Irgm3*, or the full repertoire of IRGM proteins, exhibit differences in disease progression during *Mtb* infection. We show that mice deficient in both *Irgm1* and *Irgm3* are not susceptible to *Mtb*, exhibiting a rescue phenotype compared to *Irgm1*-deficient mice. We also demonstrate that despite significant changes in the levels of certain disease associated cytokines in their lungs, mice deficient in all three IRGM proteins show the same level of host protection as wild type mice until almost one year following infection. Therefore, the increased susceptibility of *Irgm1*^-/-^ mice to acute pulmonary *Mtb* infections cannot be satisfyingly explained by a defect in cell-autonomous immunity, as proposed previously, but rather results from disrupted inter-regulatory relationships between functionally divergent *Irgm* isoforms.

## RESULTS

### *Irgm1*^-/-^ mice are acutely susceptible to infection with *Mtb*

To investigate the significance of IRGM proteins in host protection during *Mtb* infection, we first sought to recapitulate the established observation that *Irgm1*^-/-^ mice are highly susceptible to *Mtb* (14, 15). WT and *Irgm1*^-/-^ mice were infected with a low dose of *Mtb* strain H37Rv by the aerosol route. IFNγ receptor knockout (*IFNγR*^-/-^) mice were included as a control to represent the complete loss of downstream IFNγ signaling. Consistent with the results of previously published studies, the bacterial burden in *Irgm1*^-/-^ lungs was significantly higher (~1.2 log10 CFUs, *P* < 0.01) than WT at 5 weeks post-infection (Fig. 1A). Similarly, the bacterial burden in the spleens of *Irgm1*^-/-^ mice was increased (~1 log_10_ CFUs, *P* < 0.01) relative to WT (Fig. 1B). In a separate survival experiment, mice were infected with a low dose of *Mtb* by the aerosol route. All *Irgm1*^-/-^ mice and *IFNγR*^-/-^ mice succumbed to *Mtb* infection by 6 weeks post-infection, while WT mice survived beyond 150 days post-infection (Fig. 1C). Taken together, these data corroborate previously published results and indicate that mice lacking *Irgm1* are acutely susceptible to *Mtb*, exhibiting uncontrolled bacterial burden and early death (14, 15).

**FIG 1.**
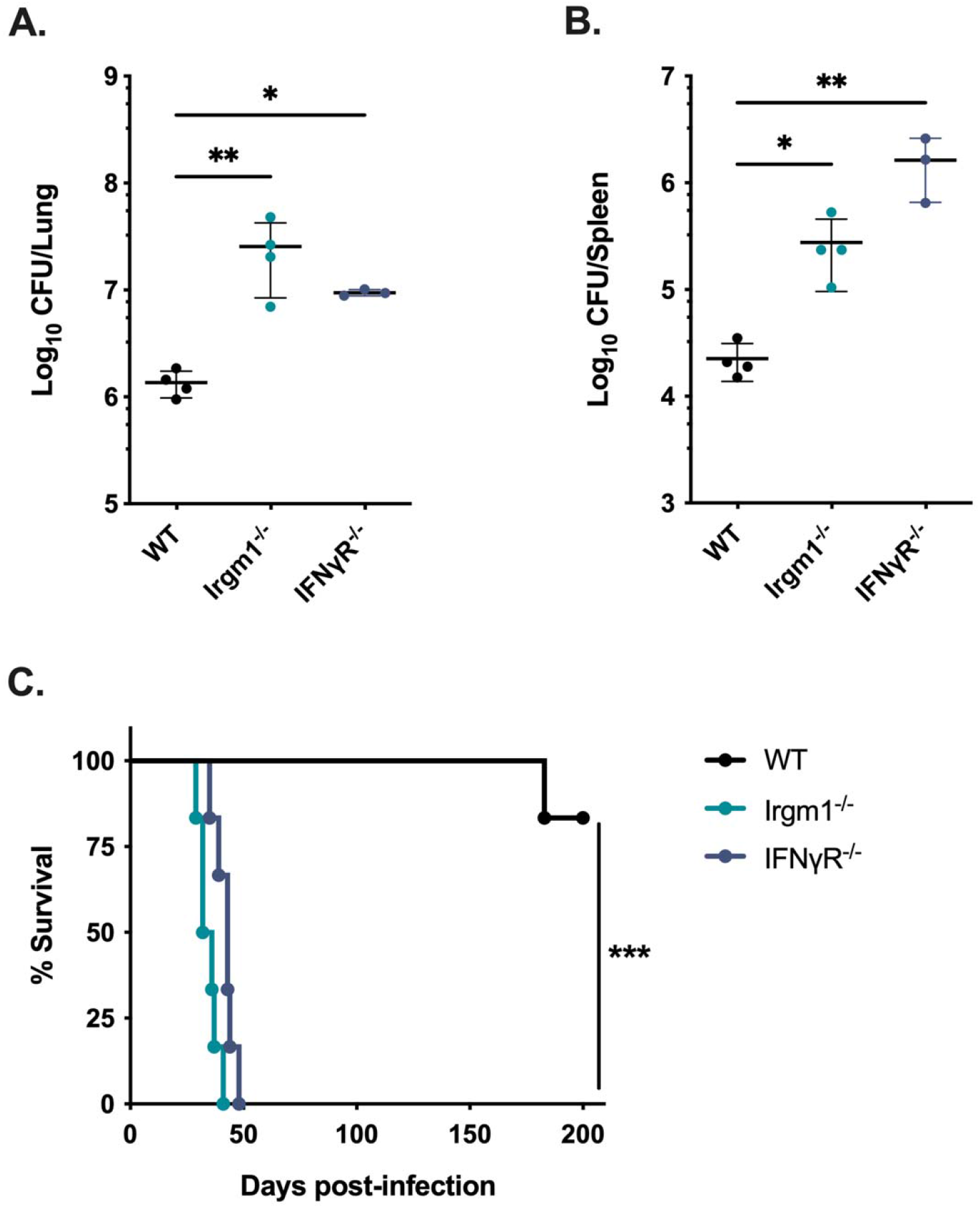
Mice lacking *Irgm1* rapidly succumb to pulmonary *Mtb* infection. WT, *Irgm1*^-/-^, and *IFNγR*^-/-^ mice were infected with *Mtb* H37Rv by the aerosol route (Day 0, 50-200 CFUs). Lungs and spleens were collected at 5 weeks post-infection and used to quantify bacterial CFUs. (A) Bacterial burden in the lungs and (B) spleens of mice. Each point represents a single mouse, data are from one experiment, with 4 male mice per group and are representative of two similar experiments. Statistics were determined via Kruskal-Wallis test by ranks and Dunn’s multiple comparisons test (* *P* < 0.05, ** *P* < 0.01). (C) WT, *Irgm1*^-/-^, and *IFNγR*^-/-^ mice were infected with *Mtb* H37Rv by the aerosol route (Day 0, 50-150 CFU) and their relative survival was quantified (Mantel-Cox test, *** *P* < 0.001). Data are from one experiment with 6 male mice per group and are representative of two similar experiments.

### Host protection against *Mtb* is restored in mice deficient in both *Irgm1* and *Irgm3*

While *Irgm1*-deficient mice are highly susceptible to *Mtb* infection, the role of the two remaining IRGM proteins (IRGM2 and IRGM3) in host protection remains unclear. Because murine IRGM proteins exhibit non-redundant functions in host protection during other infections, we tested whether knocking out *Irgm3* in an *Irgm1*^-/-^ background (*Irgm1/3*^-/-^) alters host susceptibility to *Mtb*. In contrast with *Irgm1*^-/-^ hosts, mice deficient in both *Irgm1* and *Irgm3* controlled *Mtb* burden in the lungs (Fig. 2A) and spleens (Fig. 2B) similar to WT mice at 4-weeks post-infection. Moreover, *Irgm1/3*^-/-^ mice maintained control of *Mtb* burden up to 100 days post-infection (Fig. S1A and B). Additionally, *Irgm1/3*^-/-^ mice exhibited no survival defect relative to WT mice at up to 200 days post-infection with *Mtb* (Fig. 2C). These results demonstrate that knocking out *Irgm3* in *Irgm1*-deficient mice rescues control of *Mtb* burden and restores long-term survival.

**FIG 2.**
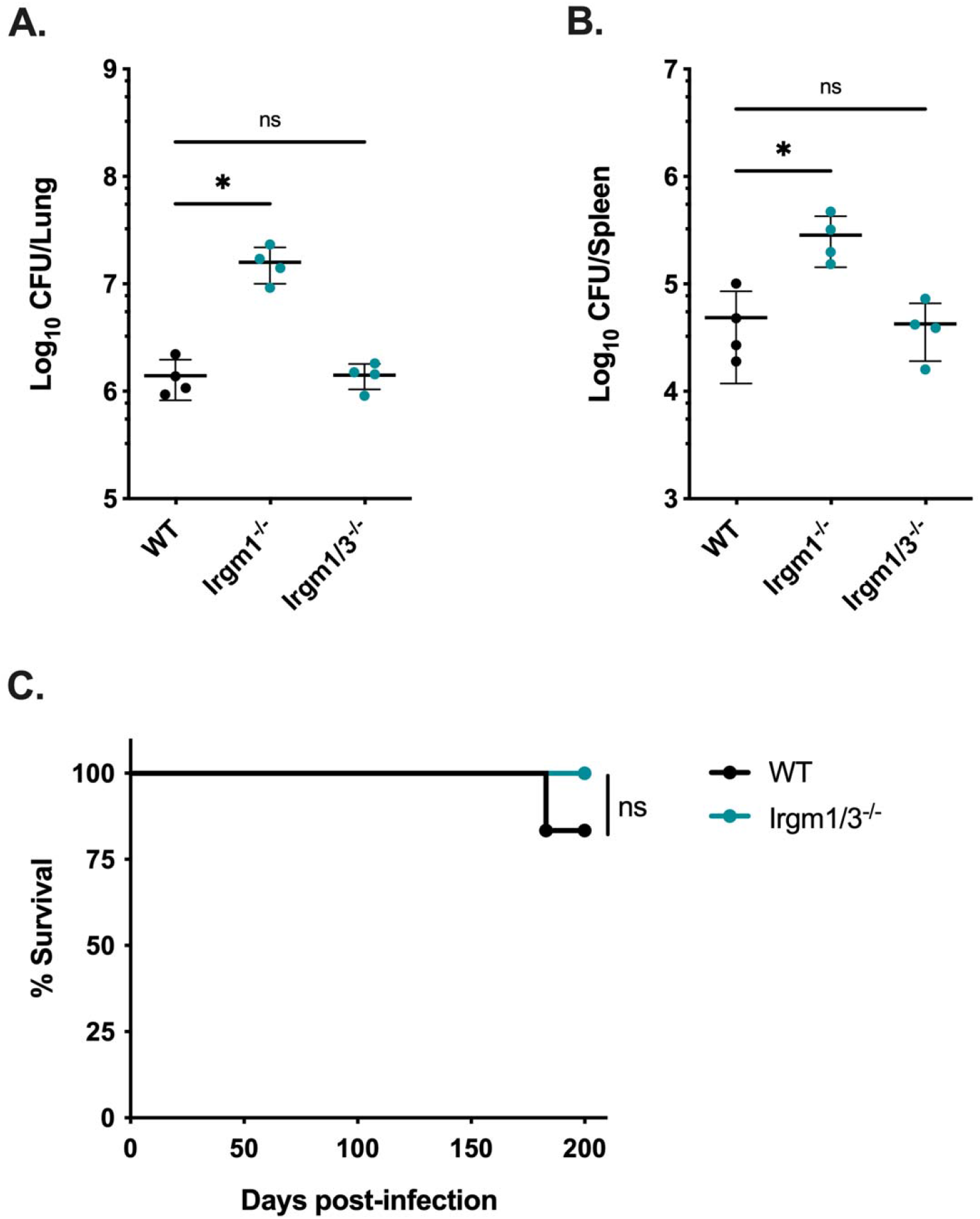
Mice deficient in both *Irgm1* and *Irgm3* are not susceptible to *Mtb* infection. WT, *Irgm1*^-/-^, and *Irgm1/3*^-/-^ mice were infected with *Mtb* H37Rv by the aerosol route (Day 0, 50-150 CFUs). Lungs and spleens were collected at 4 weeks post-infection and used to quantify bacterial CFUs. (A) Bacterial burden in the lungs and (B) spleens of mice. Each point represents a single mouse, data are from one experiment, with 4 female mice per group and are representative of 4 similar experiments. Statistics were determined via Kruskal-Wallis test by ranks and Dunn’s multiple comparisons test (* *P* < 0.05, ns = not significant). (C) WT or *Irgm1/3*^-/-^ mice were infected with *Mtb* H37Rv by the aerosol route and their relative survival was quantified (Mantel-Cox test, ns = not significant). Data are from on experiment with 6 male mice per group and are representative of 3 similar experiments.

### pan*Irgm*^-/-^ mice maintain control at one-month post-infection with *Mtb*

Although *Irgm1/3*^-/-^ mice controlled *Mtb* burden and did not have a survival defect, it remained possible that *Irgm2* is required to maintain control in the context of *Irgm1/3* deficiency. To determine whether deficiency in the entire *Irgm* locus increases susceptibility to *Mtb* after the onset of Th1 immunity, we infected pan*Irgm*^-/-^ (*Irgm1*^-/-^ *Irgm2*^-/-^ *Irgm3*^-/-^) mice with *Mtb* H37Rv by the aerosol route and compared their susceptibility to WT mice. At 4 weeks post-infection, the bacterial loads in the lungs and spleens of pan*Irgm*^-/-^ mice were not significantly different from WT mice (Fig. 3A and B). Lung sections from each group of mice were stained with hematoxylin and eosin (H&E) and used to estimate the degree of tissue damage (Fig. S2A). Consistent with the CFU data, the relative area of lung damage in WT and pan*Irgm*^-/-^ mice was comparable (Fig. 3C and D). These results contrast sharply with the phenotype of *Irgm1*^-/-^ mice following *Mtb* infection, where lung CFUs are between 1 and 2-log_10_ higher than WT and more than 70% of the air space is obstructed by lesions at 4 weeks post-infection (14) (Fig. 1). Altogether, our results suggest that the full repertoire of mouse IRGM proteins is dispensable for host resistance at one-month post-infection.

**FIG 3.**
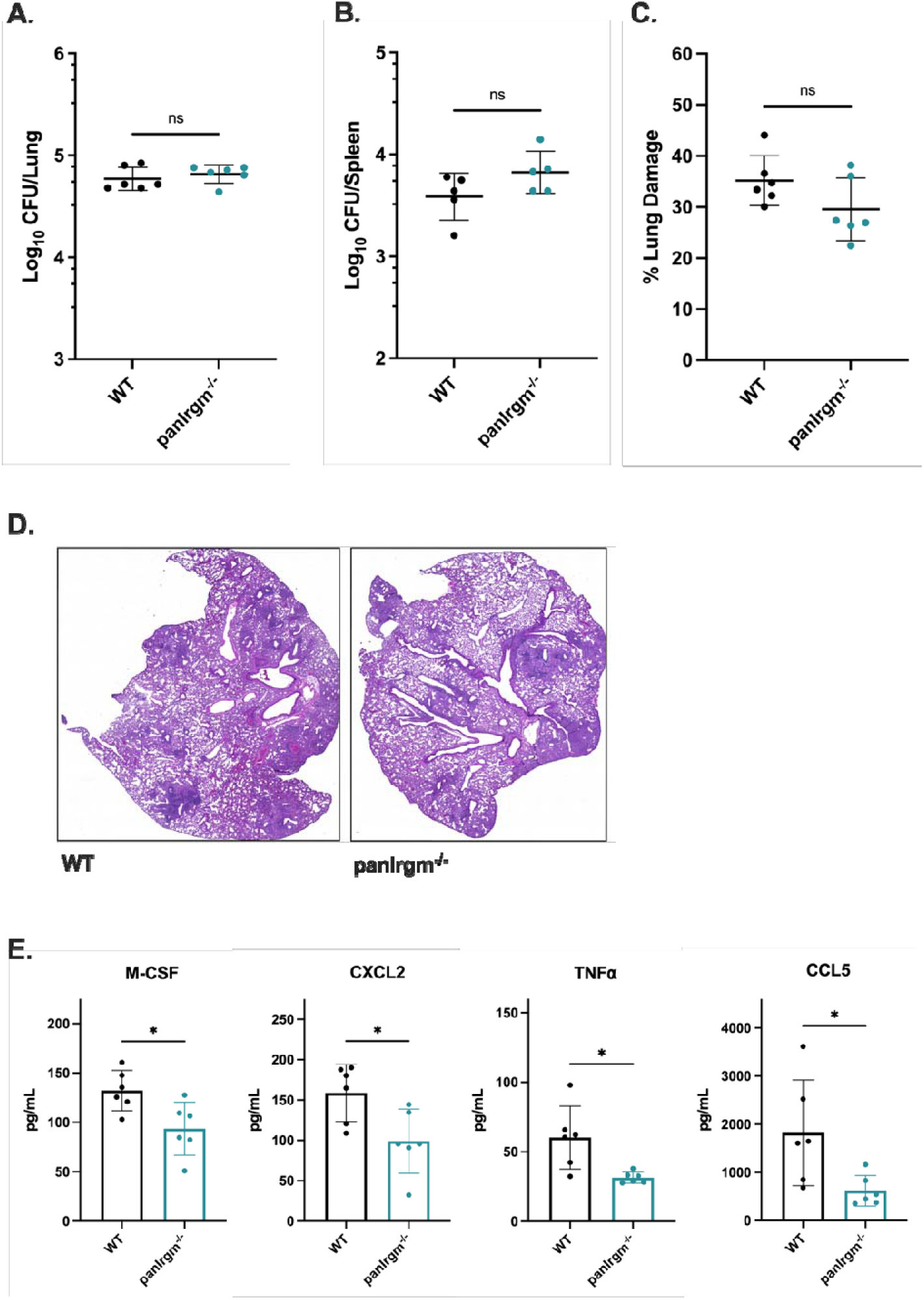
*Mtb* disease phenotypes in pan*Irgm*^-/-^ mice at 4 weeks post infection. WT or pan*Irgm*^-/-^ mice were infected with H37Rv *Mtb* by the aerosol route (~200-250 CFUs). At 4 weeks post-infection samples were collected from the lungs and spleens for phenotyping disease susceptibility and cytokines. (A) Bacterial burden in the lungs and (B) spleens of mice. (C) Relative area of damaged tissue and (D) representative images from formalin-fixed paraffin embedded lung sections that were H&E stained and used for damage quantification. (E) Concentration of cytokines (M-CSF, CXCL2, TNFͰ, or CCL5) in lung homogenates from infected mice. Each point represents a single mouse, data are from one experiment, with 4-6 male mice per group. Statistics were determined via Mann-Whitney test (* *P* < 0.05, ns = not significant).

Next, we characterized the cytokine profile in the lungs of pan*Irgm*^-/-^ mice at 4 weeks post-infection relative to WT mice. Analysis of a 32-Plex cytokine array performed on lung homogenates revealed that pan*Irgm*^-/-^ lungs contained significantly less M-CSF, CXCL2, TNFͰ and CCL5 (*P* < 0.05) than WT mice (Fig. 3E). Importantly, the levels of IFNγ were not significantly different between WT and pan*Irgm*^-/-^ lungs (Fig. S2B). Taken together, we conclude that the full repertoire of IRGM proteins in mice significantly influences a suite of cytokine responses in the lung early after the onset of adaptive immunity, but without substantially altering disease susceptibility at this stage of infection.

### pan*Irgm*^-/-^ mice exhibit higher bacterial burden and altered cytokines during late-stage *Mtb* infection

To investigate disease progression in the pan*Irgm*^-/-^ mice, we investigated disease metrics at 24 weeks post-infection. We found a slight but significant (~0.4 Log_10_; *P* < 0.05) increase in lung CFUs of pan*Irgm*^-/-^ mice relative to WT (Fig. 4A). There was no significant difference in the bacterial burden in the spleens (Fig. 4B). The relative area of damaged lung tissue was estimated from H&E-stained tissue sections, and there was no statistically significant difference between WT and pan*Irgm*^-/-^ mice (Fig. 4C and D, Fig. S3B). However, we noted that despite little variability in lung CFU numbers among the pan*Irgm*^-/-^ mice, the degree of lung tissue damage was more variable among the pan*Irgm*^-/-^ mice. Examples of both highly damaged and minimally damaged lungs relative to WT were present among pan*Irgm*^-/-^ sections (Fig. 4D).

**FIG 4.**
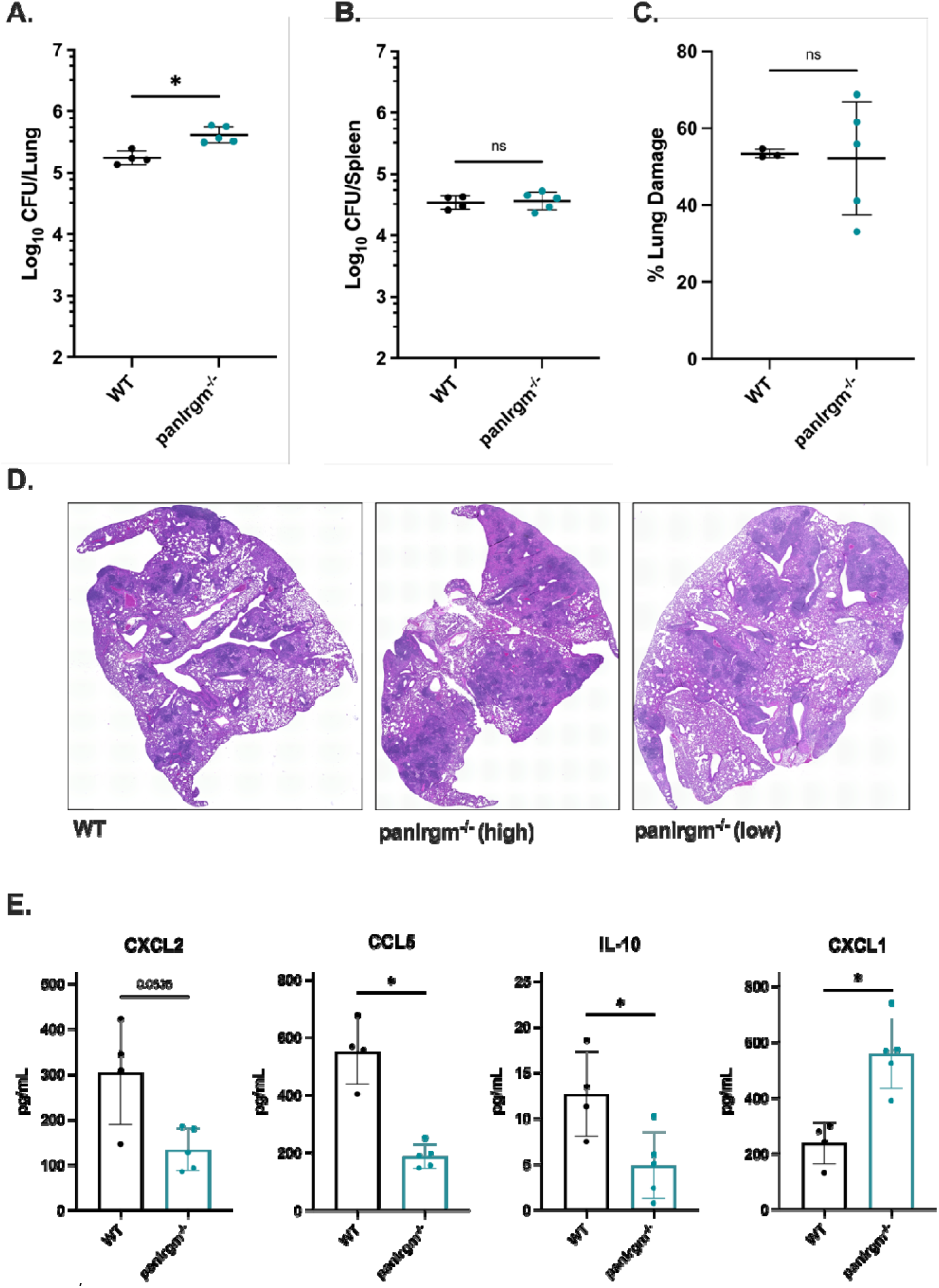
pan*Irgm*^-/-^ mice bacterial burden and cytokine response at 24 weeks post infection. WT or pan*Irgm*^-/-^ mice were infected with H37Rv *Mtb* by the aerosol route. At 24 weeks postinfection samples were collected from the lungs and spleens for phenotyping disease susceptibility. (A) Bacterial burden in the lungs and (B) spleens of mice. (C) Relative area of damaged tissue and (D) representative images from formalin-fixed paraffin embedded lung sections that were H&E stained and used for damage quantification. Examples of relatively highly damaged (“pan*Irgm*^-/-^ high”) and minimally damaged (“pan*Irgm*^-/-^ low”) lungs are shown. (E) Cytokines were quantified by multiplex ELISA. Shown are concentrations of select cytokines in lung homogenates from infected mice. Each point represents a single mouse, data are from one experiment, with 4 or 5 female mice per group. Statistics were determined via Mann-Whitney test (* *P* < 0.05, or exact *P* value shown for trends above significance threshold, ns = not significant).

Because pan*Irgm*^-/-^ mice demonstrated altered bacterial burden in the lungs at 24 weeks, we additionally investigated the cytokine response at this timepoint. Here, we observed a significant decrease in CCL5 (*P* < 0.05) and a trend toward decreased CXCL2 (*P* = 0.0635) in pan*Irgm*^-/-^ lungs (Fig. 4E). IFNγ levels were unaffected by the loss of all three IRGM proteins (Fig. S3A). When we analyzed the remaining cytokines in the 32-Plex panel, we identified several that were associated with the loss of IRGM proteins at the 24-week timepoint. Specifically, IL-10 was significantly (*P* < 0.05) decreased in pan*Irgm*^-/-^ mice relative to WT mice, while CXCL1 was significantly (*P* < 0.05) increased (Fig. 4E). Overall, our results indicate the loss of all *Irgm* genes shifts the abundance of select cytokines in the lungs of mice during chronic disease.

### pan*Irgm*^-/-^ mice show impaired survival to infection

Considering pan*Irgm*^-/-^ mice showed elevated lung bacterial burden and altered cytokines at 24 weeks post-infection, we investigated the overall survival of WT and pan*Irgm*^-/-^ mice over the course of infection. pan*Irgm*^-/-^ mice began to die earlier (*P* < 0.01), with less than a 40% chance of survival by 52 weeks post-infection compared to ~90% survival in WT mice (Fig.5). Thus, the loss of all IRGM proteins impacts overall disease susceptibility to *Mtb* but does so very late in the course of disease in mice.

**FIG 5.**
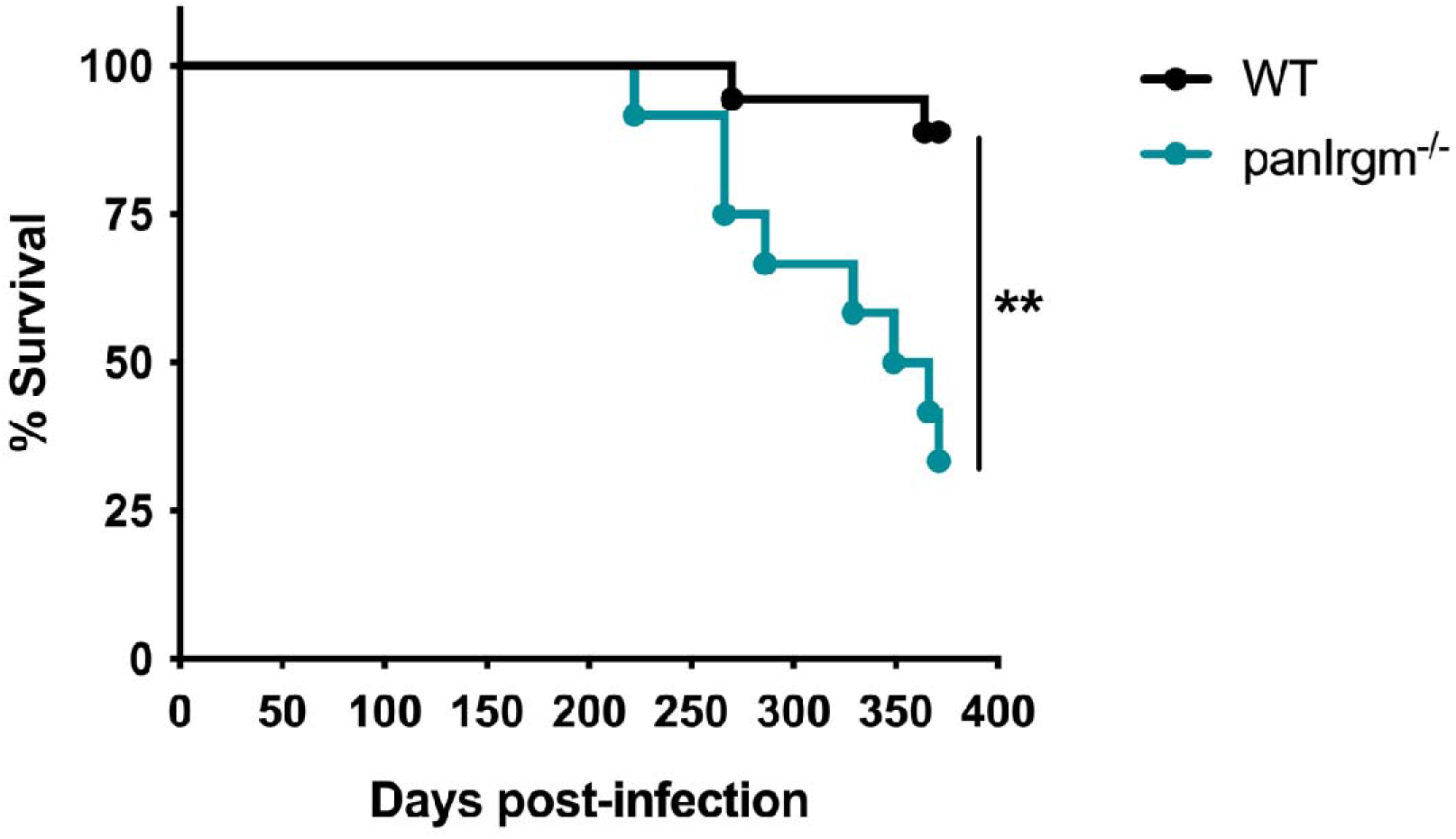
pan*Irgm*^-/-^ mice survival phenotype after long-term *Mtb* infection. WT or pan*Irgm*^-/-^ mice were infected with H37Rv *Mtb* by the aerosol route (~200-250 CFUs; as per Figure 4). Relative survival of WT and pan*Irgm*^-/-^ mice following *Mtb* infection by the aerosol route (Mantel-Cox test, ** *P* < 0.01). Data are from one experiment with 12-18 male mice per group.

## DISCUSSION

Cell-intrinsic immunity is a fundamental mechanism through which hosts defend against a variety of pathogens, including viruses, bacteria, and parasites. IFNγ is vital in coordinating cell-autonomous immune responses, yet the context-dependent mechanisms by which downstream IRGs facilitate host protection are incompletely understood. Comparing the functions of human and murine IRGM proteins in the context of different pathogens and during autoimmune disease continues to provide new insights into the ways these proteins regulate IFNγ-dependent immune responses (26). In this work, we examined whether functional relationships between *Irgm1, Irgm2*, and *Irgm3* genes significantly influence host protection in the context of early (4 weeks) and late (24 weeks) stages of *Mtb* infection in mice. Our results suggest that a balance between IRGM proteins is required to effectively protect against TB.

Similar to previous studies, we found that *Irgm1*^-/-^ mice succumb to *Mtb* infection rapidly with uncontrolled bacterial replication (14, 15). Various interrelated explanations for the extreme susceptibility of *Irgm1*^-/-^ mice to *Mtb* have been proposed, including defective phagosome maturation, impaired autophagosome biogenesis and/or delivery of mycobacteria to late endosome/lysosome compartments, and dysregulated T cell survival (5, 8, 14, 15). However, the contribution of each of these mechanisms to the loss of host protection remains a matter of debate and is dependent on the pathogen. For example, *Irgm1*^-/-^ mice are also susceptible to *S. typhimurium, L. monocytogenes, C. trachomatis*, and *T. gondii* infection (26). During *T. gondii* infection, *Irgm1* contributes to cell-autonomous control of the pathogen by regulating activation of GKS IRG proteins and their accumulation on the parasitophorous vacuole membrane, ultimately coordinating destruction of the pathogen (27). In the context of mycobacterial and listerial infections, it was initially proposed that IRGM1 is critical for phagocytosis, directly localizing to the phagosome membrane to promote maturation (14, 28, 29). However, experiments using improved antibodies and more extensive controls to dissect IRGM1 localization during mycobacterial and listerial infection *in vitro* could not confirm that IRGM1 directly associates with the phagosomal membrane (30).

In the context of both IRGM1 and IRGM3, studies examining the simultaneous loss of both proteins discovered that the balance of different IRGM proteins modulates disease outcome in a pathogen-dependent manner. While the susceptibility of *Irgm1*^-/-^ mice to *S. typhimurium* is reversed in *Irgm1/3*^-/-^ mice, the susceptibility remains during infection with *C. trachomatis* and *T. gondii*, suggesting distinct mechanisms of protection that are dependent on the pathogen (23, 24). We observed that at the onset of adaptive immunity, *Mtb* infection resembles *S. typhimurium*, with the rapid disease progression of *Irgm1*^-/-^ mice being entirely reversed in *Irgm1/3*^-/-^ mice that survive far into the chronic phase of disease with no changes in bacterial control. *Mtb* infection perturbs mitochondrial functions in macrophages, and even in uninfected *Irgm1*-deficient mice macrophages exhibit defective mitochondrial quality control (31, 32). Taken together, this suggests that rather than directly controlling *Mtb* replication in macrophages, a balance between IRGM1 and IRGM3 may be required to maintain disease tolerance by regulating mitochondrial functions, and that loss of this balance is the source of susceptibility to *Mtb* in *Irgm*^-/-^ mice.

How IRGM1 and IRGM3 together regulate the overall host response during *Mtb* infection is not entirely clear and may be multifactorial. Taken together, past studies indicate that IRGM functions overlap with both autophagy and type I IFN pathways; importantly both of these are previously proposed correlates of TB disease progression (32–35). Specifically, previous studies indicate that the loss of murine *Irgm1* or human *IRGM* is consistently associated with defects in autophagy during infection. This autophagy dysfunction drives a shift toward pro-inflammatory metabolism, increased inflammasome activation, and death of proliferating T cells that results in lymphopenia (36, 37). As mentioned above, the loss of *Irgm1* also affects mitochondrial quality control, which leads to exacerbated type I IFN responses in macrophages (32). Our results highlight the importance of *Irgm* balance in controlling protective responses, independent of bacterial replication, but the underlying mechanisms driving unbalanced *Irgm* responses remain to be understood. Given the complex roles of type I IFNs in TB disease (6, 34), it will be important to confirm how the balance of *Irgm1* and *Irgm3* in mice affects type I IFN responses during Mtb infection, and whether this is related to mis-targeting of GKS proteins in immune cells (32). Although detailed mechanistic analysis suggests that IRGM1 does not directly localize to the Mycobacterial containing vacuole and is not likely a direct effector at the phagosome, it remains possible that the loss of *Irgm1* leads to increased IRGM3 activity and dysfunction in maintaining stability of intracellular membrane compartments (30).

Given the clear genetic interactions between *Irgm1* and *Irgm3* during *Mtb* infection it was also important to consider the role *Irgm2* plays in TB susceptibility using a mouse lacking all three IRGM proteins (pan*Irgm*^-/-^). We observed that pan*Irgm*^-/-^ mice showed no changes to bacterial burden or lung damage early after the initiation of adaptive immunity. However, there were some significant differences in cytokine production in the lungs. At a much later stage of disease, pan*Irgm*^-/-^ mice displayed marginally increased bacterial burden in the lungs, more variable tissue damage, and ultimately died earlier than wild type animals. The late presentation of a survival defect in pan*Irgm*^-/-^ mice, following over 300 days of infection, further suggests that IRGM proteins do not strongly contribute to direct control of *Mtb* replication as proposed previously. Instead, it is more likely that the IRGM proteins are involved in controlling the inflammatory environment and tissue damage in the lungs, which appears to be critically important during very late stages of infection, or advanced host age. Of the cytokines examined, we consistently observed changes in chemokines that drive immune cell recruitment into the infected lung environment, including CXCL1 that is a known correlate of TB disease severity in diverse mice and humans (38–43).

The late emergence of a survival defect in pan*Irgm*^-/-^ mice during *Mtb* infection contrasts sharply with *T. gondii* infection, where pan*Irgm*^-/-^ mice survive poorly, at a frequency only slightly higher than *Irgm1*^-/-^ or *Irgm1/3*^-/-^ mice (23, 25). Our data clearly indicate a need for IRGM proteins very late during infection. Age is a factor known to contribute to TB disease susceptibility, and changes during aging, including lower nutrition and immunosuppression or immune dysregulation, function via pleotropic mechanisms (44, 45). Exemplifying the phenomenon of “inflammaging”, IL-12, TNFͰ and IL-1β increase in the lung as the host ages, yet cause and effect mechanisms for this are difficult to isolate. Whether IRGM proteins are required to maintain lung protection, perhaps contributing to the regulation of inflammatory responses, as the host ages is an important hypothesis to consider. How our results relate to human IRGM functions also remains to be determined. For example, it is unclear whether *IRGM* polymorphisms that have been identified as correlates of TB control or progression alter cell-autonomous control of *Mtb* in human macrophages, and/or have pleiotropic effects on the immune response to *Mtb* via autophagy and metabolism.

In conclusion, in this work we have begun to address the need for a balance of IRGM proteins during *Mtb* infection in mice for long-term protection. *Irgm1* is necessary for early control of *Mtb* infection in mice when it is lost individually, but the imbalance created by the loss of *Irgm1* can be repaired by eliminating *Irgm3* or all IRGM proteins. However, as the infection progresses to a very late stage in an aging host, the loss of all IRGM proteins becomes detrimental to host survival and is associated with early death.

## MATERIALS AND METHODS

### Ethics statement

Mouse studies were performed in strict accordance using the recommendations from the Guide for the Care and Use of Laboratory Animals of the National Institute of Health and the Office of Laboratory Animal Welfare. Mouse studies were performed using protocols approved by the Institutional Animal Care and Use Committee (IACUC) for each institution, in a manner designed to minimize pain and suffering in *Mtb*-infected animals. IACUC numbers for each institution includes the University of Massachusetts Medical School (A3306-01); Duke University (A221-20-11); Michigan State (PROTO202200127). Any animal that exhibited severe disease signs was immediately euthanized in accordance with IACUC approved endpoints.

### Mouse strains and infection with *Mtb*

WT C57BL/6J mice were purchased from The Jackson Laboratory (#000664). pan*Irgm*^-/-^ mice were generated by the lab of Dr. Jörn Coers at Duke University as described previously (46). All mice were housed in a specific pathogen-free facility under standard conditions (12hr light/dark, food and water *ad libitum*). Mice were infected with *Mtb* between 8-12 weeks of age with the H37Rv strain of *Mtb* (PDIM positive). For aerosol infections, *Mtb* was cultured in 7H9 media supplemented with oleic acid-albumin-dextrose-catalase OADC enrichment (Middlebrook) and 0.05% Tween 80 (Fisher). Prior to all *in vivo* infections, *Mtb* cultures were washed, resuspended in phosphate-buffered saline (PBS) containing 0.05% Tween 80, and sonicated or filtered through a 40 μM filter to generate a single-cell suspension. For infections of WT, *Irgm1*^-/-^ and *Irgm1/3*^-/-^ mice, an inoculum between 50-200 CFUs was delivered by an aerosol generating Glas-Col chamber. For WT versus pan*Irgm*^-/-^ infections, an inoculum of ~200-250 CFUs was delivered by an aerosol generating Madison chamber (University of Wisconsin at Madison) to the groups of mice as indicated. To determine the inoculation dose, 5 mice were euthanized at 1 day post-infection and CFUs were enumerated from lung homogenates, as described below.

### Bacterial burden quantification

At 1 day, 4 weeks, and 24 weeks post-infection, mice were euthanized by overdose with isoflurane (Covetrus) and the spleens and lungs were removed aseptically. For enumeration of viable bacteria, organs were individually homogenized in PBS-Tween 80 (0.05%) by bead beating (MP Biomedical), and 10-fold dilutions were plated on 7H10 agar (Middlebrook) plates containing OADC enrichment (Middlebrook) and 50 μg/mL Carbenicillin, 10μg/mL Amphotericin B, 25 μg/mL Polymyxin B, and 20 μg/mL Trimethoprim (Sigma). Plates were incubated at 37°C for 3-4 weeks and individual colonies were enumerated to calculate CFUs.

### Lung pathology and estimation of damage

In parallel with bacterial burden quantification, one lung lobe from each mouse was reserved for histology and fixed in 10% neutral buffered formalin. Lungs were submitted to the Duke University Pathology core facility where they were paraffin embedded, sectioned at 5 μM, and stained with hematoxylin and eosin. Lung sections were imaged at 10X magnification and stitched into whole-lung images for each mouse (Keyence). The relative area of damaged tissue was estimated using QuPath v0.3.2 (47). For this, a pixel classifier was trained to identify damaged versus undamaged areas of H&E stained mouse lung sections, based on 132 damaged annotations and 96 undamaged annotations across 11 training images, which were imported to train the ANN_MLP classifier at moderate resolution. The quality of the classifier was visually inspected across diverse samples by overlaying live annotation predictions with each H&E image. The resulting pixel classifier was loaded into QuPath and used to estimate the damaged area of each lung sample, relative to the total lung area (Fig. S1 and S2).

### Quantification of cytokines in tissue homogenates

Murine lung homogenates were centrifuged to remove cellular debris, and the supernatants were filtered through 0.2 μM filters. 32 cytokines/chemokines were quantified via a Discovery Assay (Eve Technology; MD31). Observed concentrations were used for all downstream analyses.

### Statistical analysis

Statistical analyses were performed using Prism 9 (Graph Pad) software. Bacterial burden, lung damage, and cytokine differences between two groups were analyzed using Mann-Whitney test. When more than two groups were compared, Kruskal-Wallis test by ranks and Dunn’s multiple comparisons test were selected. Differences in survival were graphed using Kaplan-Meier curves, and statistical significance was assigned via Mantel-Cox testing. Throughout, *P* value thresholds are noted as ns = not significant, * *P* < 0.05, ** *P* < 0.01, *** *P* < 0.001.

## SUPPLEMENTAL MATERIAL

Figure S1-S3, PDF file

## ACKNOWLEDGEMENTS

We thank Dr Greg Taylor for insightful manuscript comments, Summer Harris and Emily Hunt for technical assistance and members of the Olive, Smith, Sassetti and Coers labs for helpful feedback. This work was funded by National Institutes of Health grants AI148243 and AI103197 to JC; AI132130 to C.M. Sassetti; AI165618 to A.O; a Whitehead Scholar Award and an NIH Director’s New Innovator Award (GM146458) to C.M. Smith. Biocontainment work was partially performed in the Duke Regional Biocontainment Laboratory, which received partial support for construction from the National Institutes of Health, National Institute of Allergy and Infectious Diseases (UC6-AI058607; G20-AI167200).

