## Supplemental Figures S1-S3 for "Differential requirement for IRGM proteins during tuberculosis infection in mice"

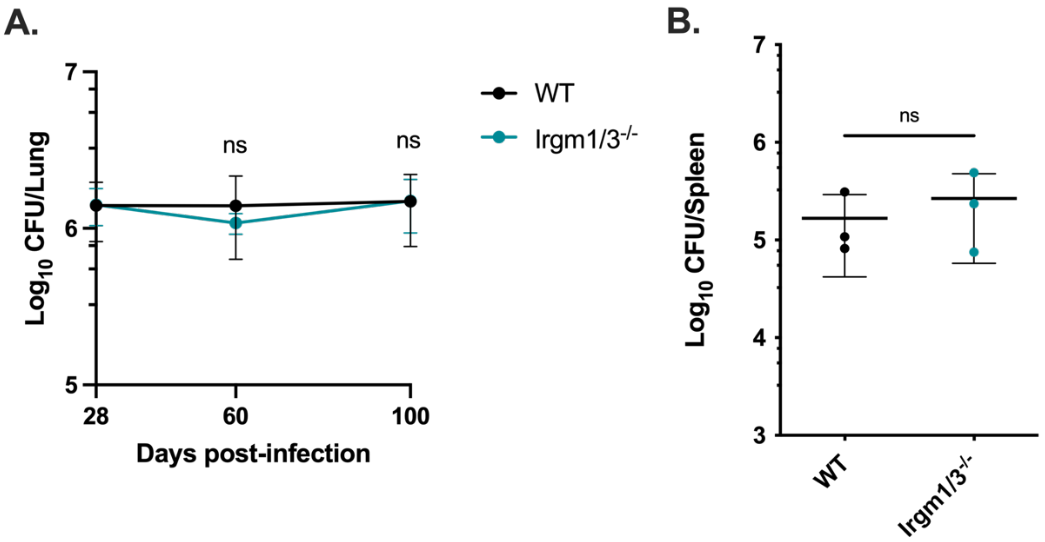


**FIG S1. *Irgm1/3$^{-/-}$* mice control *Mtb* burden during long-term infection**. Following aerosol infection with *Mtb* H37Rv (Day 0 dose of 50-150 CFUs), CFUs were quantified from the lungs (A) and spleens (B) of mice at 60- and 100-days post-infection (A) or 100 days post-infection (B). Each point represents a single mouse, data are from one experiment, with 3-4 female mice per group. Statistics were determined via Mann-Whitney test (ns = not significant).

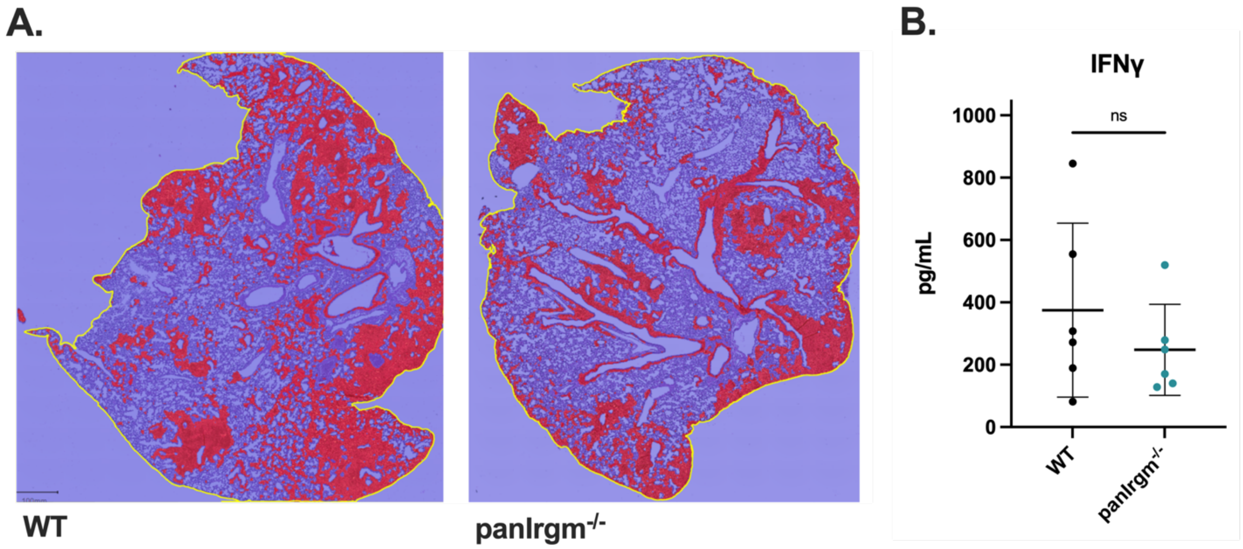


**FIG S2.** (A) QuPath pixel classifier estimates of relative damaged area in representative H&E stained lung sections from mice, collected at 4 weeks post-infection. (B) IFNγ cytokine concentration in lung homogenates from infected mice at 4 weeks post-infection. Statistics were determined via Mann-Whitney test (ns = not significant).

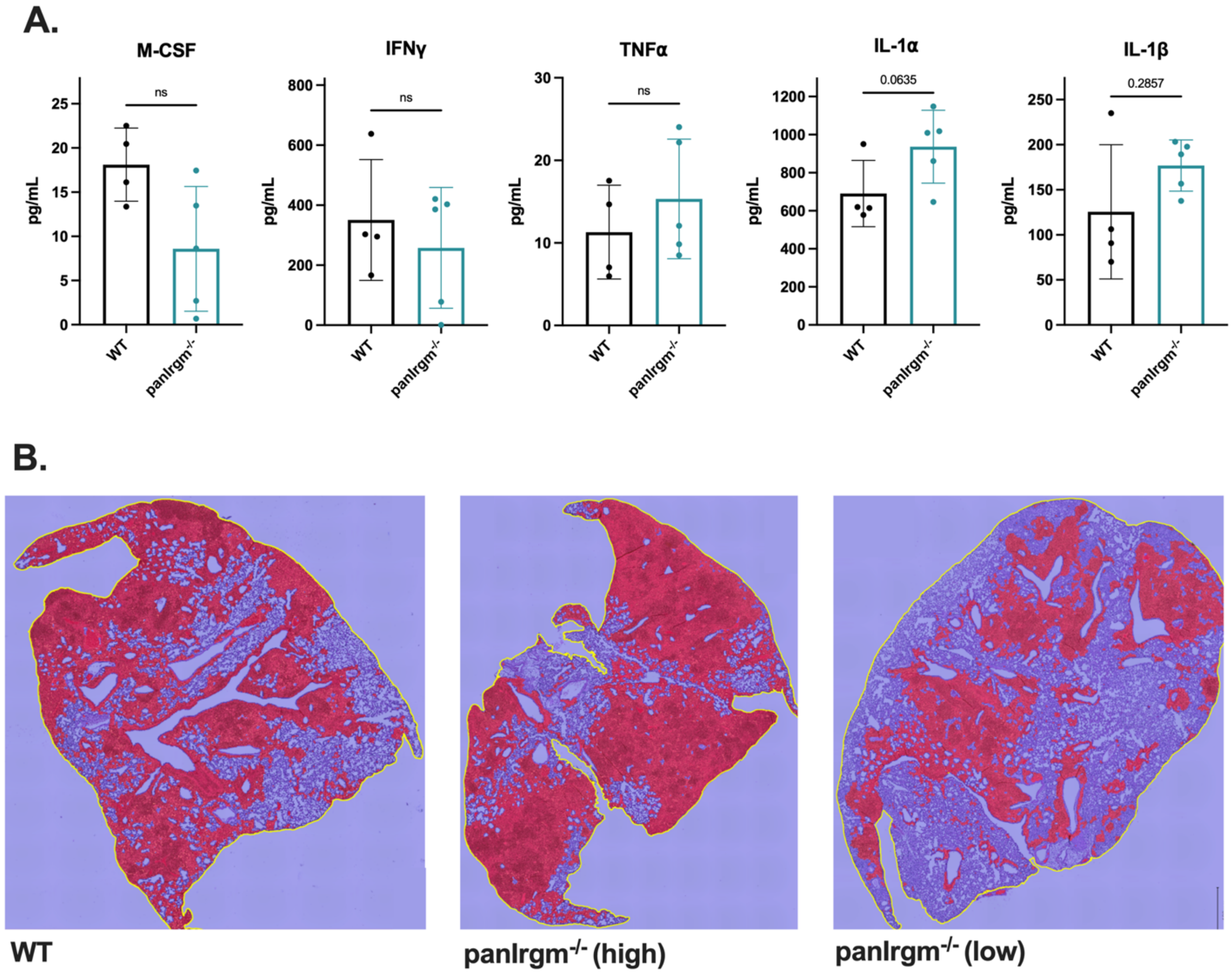


**FIG S3**. (A) Concentration of cytokines in lung homogenates from infected female mice at 24 weeks post-infection. Statistics were determined by Mann-Whitney test (ns = not significant, or exact *P* value shown for trends above significance threshold). (B) QuPath pixel classifier estimates of relative damaged area in representative H&E-stained lung sections from mice, collected at 24 weeks post-infection.
